# First versatile reverse genetics system for feline coronavirus

**DOI:** 10.1101/2024.10.02.616382

**Authors:** Izumi Kida, Tomokazu Tamura, Yudai Kuroda, Takasuke Fukuhara, Ken Maeda, Keita Matsuno

## Abstract

Feline infectious peritonitis (FIP) is a fatal disease caused by feline coronavirus (FCoV). Although multiple gene mutations in FCoV likely account for FIP pathogenesis, molecular studies for FCoV have been limited due to the lack of a suitable reverse genetics system. In the present study, we established a rapid PCR-based system to generate recombinant FCoV using the circular polymerase extension reaction (CPER) method for both serotype 1 and 2 viruses. Recombinant FCoV was successfully rescued at sufficient titers to propagate the progeny viruses with high sequence accuracy. The growth kinetics of recombinant FCoV were comparable to those of the parental viruses. We successfully generated recombinants harboring spike gene from a different FCoV strain or a reporter HiBiT-tag using the CPER method. The chimeric virus demonstrated similar characteristics with the parental virus of S gene. The reporter tag stably expressed after five serial passages in the susceptible cells, and the reporter virus could be applied to evaluate the sensitivity of antiviral inhibitors using the luciferase assay system to detect HiBiT tag. Taken together, our versatile reverse genetics system for FCoV shown herein is a robust tool to characterize viral genes even without virus isolation and to investigate the molecular mechanisms of the proliferation and pathogenicity of FCoV.

**Importance:** Feline infectious peritonitis is a highly fatal disease in cats caused by feline coronavirus variants that can infect systemically. Because of a lack of versatile toolbox of manipulating the feline coronavirus genome, an efficient method is urgently needed for studying virus proteins responsible for the severe disease. Herein, we established a rapid reverse genetics system for the virus and demonstrated the capability of the recombinant viruses to be introduced desired modifications or reporter genes without any negative impacts on virus characteristics in cell culture. Recombinant viruses are also useful to evaluate antiviral efficacy. Overall, our system can be a promising tool to reveal the molecular mechanisms of viral life cycle of feline coronavirus and disease progression of feline infectious peritonitis.

## Introduction

Feline infectious peritonitis (FIP) is a fatal disease caused by feline coronavirus (FCoV). FCoV infection is ubiquitous and usually causes mild or no symptoms in a cat population, affecting up to 90% of cats in multi-cat households (1). FCoV can be classified into two pathotypes, feline enteric coronavirus (FECV), which exhibits low pathogenicity typically causing asymptomatic or mild enteritis, and highly pathogenic FIP virus (FIPV) responsible for FIP, which originates from FECV mutated in the infected cats (2, 3). The incidence of FCoV-infected cats that succumb to FIP is estimated to be 5-12% (4, 5). There are two serotypes of FCoV: serotype-1 (FCoV-1) is the most dominant and unique to cats (6, 7), while serotype-2 (FCoV-2) is less common and emerged by recombination between serotype-2 canine coronavirus (CCoV-2) and FCoV-1 (8–11).

Pathotype switch from FECV to FIPV is linked to several gene mutations, including those in the ORF3c, ORF7b and spike (S) genes (2, 12–15). The intact forms of ORF3c and ORF7b are thought to be essential for FECV replication, but not for FIPV, suggesting these viral proteins are not directly associated with the pathotype switch (12, 13). The S protein of some coronaviruses, including FCoV-1, consists of the S1 and S2 subunits which are involved in receptor binding and recognition, immune evasion, and membrane fusion (16–18). The S protein encodes a polybasic cleavage site at the S1/S2 boundary, recognized by the host furin protease, and required for the cleavage of the S1 and S2 subunits. Cleavage-deficient mutation(s) at this site should be the most significant feature of FIPV compared to FECV (14, 15, 19–22). However, the molecular function of the amino acid changes has not been investigated using the infectious viruses due to the lack of suitable reverse genetics systems for FCoV. To understand the functions of each mutation and the molecular mechanism of the pathotype switch from FECV to FIPV, it is essential to generate recombinant virus with specific mutations and examine the virological characteristics of FCoV variants. Thus, a simple and effective reverse genetics system is urgently needed for further molecular studies of FCoV.

Infectious clones carrying the full-length FCoV cDNA under appropriate promoters have been established based on different backbones; bacterial artificial chromosomes (BAC) vector (23), vaccinia virus vector (24–26), and transformation-associated recombination (TAR) system in yeast (27), while these approaches may be difficult to rapidly modify viral genes. Recently, a PCR-based rapid assembly of infectious cDNA using circular polymerase extension reaction (CPER) has been widely applied to various RNA viruses, including SARS-CoV-2 (28–31). As a proof-of-concept study, we applied CPER method to generate infectious clones of the virulent FCoV-1 isolate of C3663 (32, 33) and FCoV-2 isolate of WSU 79-1146 (34, 35) possessing reporter genes or chimeric S genes and compared the virological features of the recombinants to the parental FCoVs.

## Result

### Establishment of CPER-based reverse genetics for FCoV

To generate recombinant FCoVs (rFCoVs) using CPER, we used a serotype-1 FCoV strain C3663 and a serotype-2 FCoV strain WSU 79-1146 as parental viruses. A total of 10 gene fragments (F1 to F10) covering the entire genome of each FCoV and an untranslated region (UTR) linker fragment encoding sequences of 18 nt of the 3’UTR of each FCoV, hepatitis delta virus ribozyme (HDVr), bovine growth hormone (BGH) polyA signal, cytomegalovirus (CMV) promotor, and 70 nt of the 5’UTR of each FCoV were cloned into plasmids. Each fragment (F1-F10 and the UTR linker) contained overlapping sequences of at least 13 nt with each other and was amplified using specific primers followed by CPER assembly (Fig. 1A). The CPER products were transfected into Fcwf-4 cells co-cultured with HEK293T cells to facilitate viral recovery (36).

**Fig 1.**
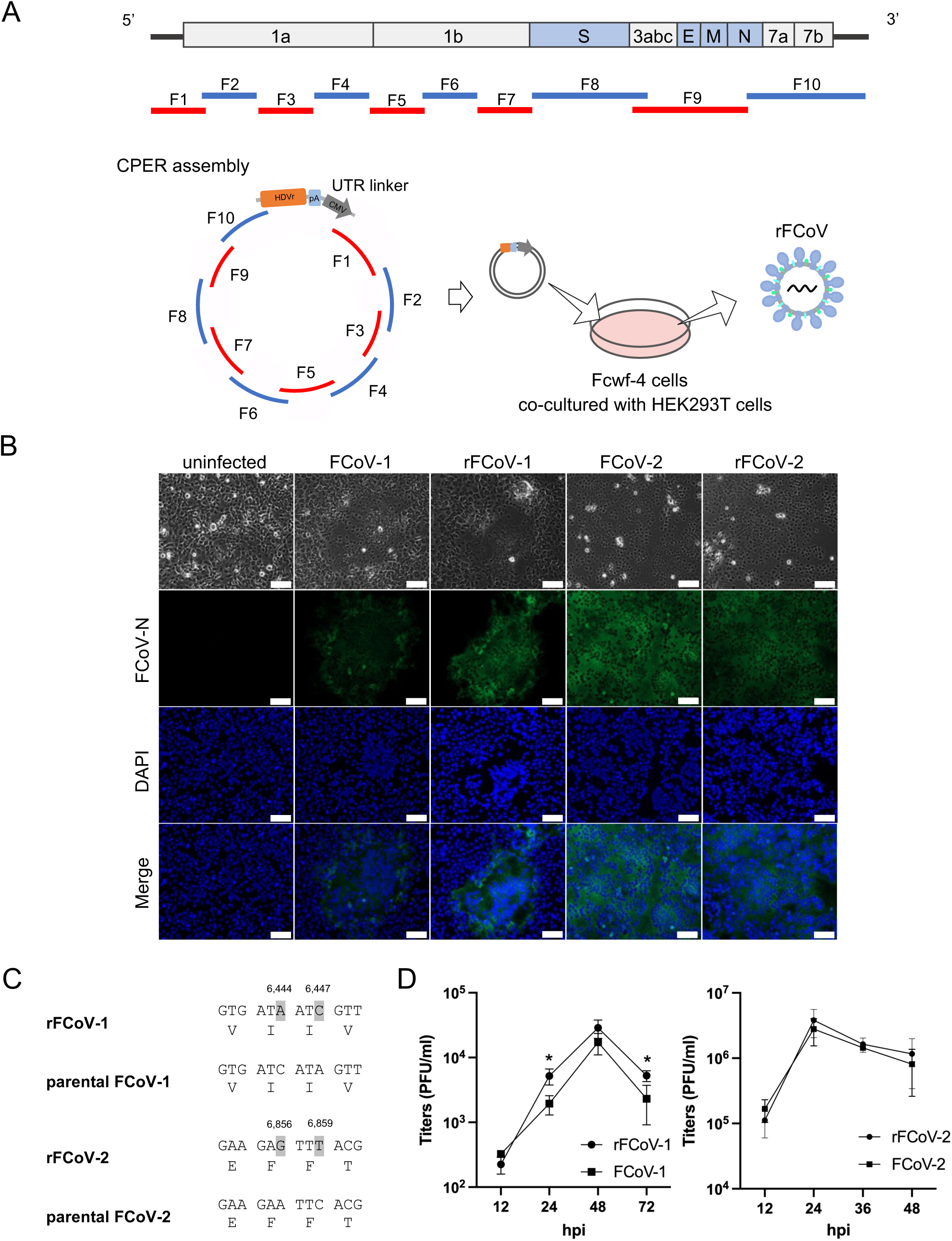
Establishment of reverse genetics for FCoV by circular polymerase extension reaction (CPER) (A) Schematic representation of recombinant virus rescue. FCoV genome (top) was divided into 10 cDNA fragments (F1-F10) covering the full-length of FCoV genome with 13 to 236 nt overlapping ends (middle) and used for a CPER assembly for generating recombinant virus (rFCoV) (bottom). The F1 to F10 fragments were assembled with an untranslated region (UTR) linker fragment by CPER and then the resulting CPER products were transfected into Fcwf-4 cells co-cultured with HEK293T cells. (B) Immunofluorescence assay of Fcwf-4 cells infected with parental FCoVs and CPER-generated rFCoVs. Viral antigen was visualized by staining with anti-FCoV N protein monoclonal antibody (green). Nuclei were stained with DAPI (blue); scale bar: 100 µm. (C) Genetic markers, 2 silent mutations (i.e. C6444A and A6447C for FCoV-1 and A6856G and C6859T for FCoV-2) were introduced into the plasmids for the corresponding fragments and confirmed in the recombinant FCoV genomes. (D) Growth kinetics of rFCoVs and parental FCoVs. Fcwf-4 cells were infected with the viruses at an MOI of 0.01, and the virus titers in the supernatants were measured from 12 to 48 or 72 hours post-infection (hpi). The presented data were expressed as mean ± SD of triplicate samples. *: *p* < 0.05 by two-tailed Student’s *t* test without adjustment for multiple comparisons.

The culture supernatants of transfected cells were harvested at 4 days post-transfection (dpt) for rFCoV-1 and at 2 dpt for rFCoV-2, when cytopathic effect (CPE) was observed, and then passaged in naïve Fcwf-4 cells. CPE was observed at 3 days post-infection (dpi) for rFCoV-1 and at 1 dpi for rFCoV-2 in the first passage (P1), and virus replication was confirmed by expression of viral antigen in the P1 cells (Fig. 1B). The infectious titers of the P1 rFCoV-1 and rFCoV-2 were 10^4.5^ PFU/ml and 10^6.5^ PFU/ml, respectively, demonstrating that rFCoVs were recovered at sufficient titers for downstream assays and capable of propagating the progeny viruses. To verify the full-length genome of P1 viruses, RNA extracted from rFCoVs was subjected to RNA-sequencing. Sequence analysis of the viruses revealed that there was no sequence difference above 35% cut-off between rFCoVs and the parental viruses except for the genetic markers (Fig. 1C). Then, we compared the growth kinetics of rFCoVs with artificially introduced parental FCoVs. The P1 viruses and parental FCoVs were infected into Fcwf-4 cells at a multiplicity of infection (MOI) of 0.01 and replicated with comparable kinetics to each other (Fig. 1D). These results suggest that the virological characteristics of rFCoVs *in vitro* are similar to those of the parental viruses.

### Construction of chimeric FCoV-2 encoding spike gene from FECV-2

To examine whether the CPER method can be applied to generate recombinant FCoVs with the desired modifications, we generated rFCoV-2 carrying FECV S proteins of WSU 79-1683, which can be responsible for pathotype switch of FCoV (37) (Fig. 2A). In the present study, we replaced S gene of FECV-2 into the F8 plasmid of FCoV-2 and subjected it to CPER with other fragments for FCoV-2 to rescue rFCoV-2 carrying FECV S protein (rFCoV-2 1683-S). The culture supernatants were harvested when CPE was observed at 3 dpt and then passaged in naïve Fcwf-4 cells. CPE was observed at 1 dpi in the P1 Fcwf-4 cells, and rFCoV-2 1683-S was recovered from the supernatant. We confirmed the replacement of S gene in rFCoV-2 1683-S by RNA-sequencing analysis. Since plaque size is different between FCoV-2 pathotypes (38) and should be governed by coronavirus S protein (39), we compared the plaque size in Fcwf-4 cells infected with rFCoV-2 1683-S and parental viruses including FCoV-2, rFCoV-2 and FECV-2 WSU 79-1683 (Fig. 2B). The plaque size of rFCoV-2 1683-S was smaller compared with those of FCoV-2 and rFCoV-2, both exhibiting FIPV phenotype, but similar to FECV-2, from which S gene was derived.

**Fig 2.**
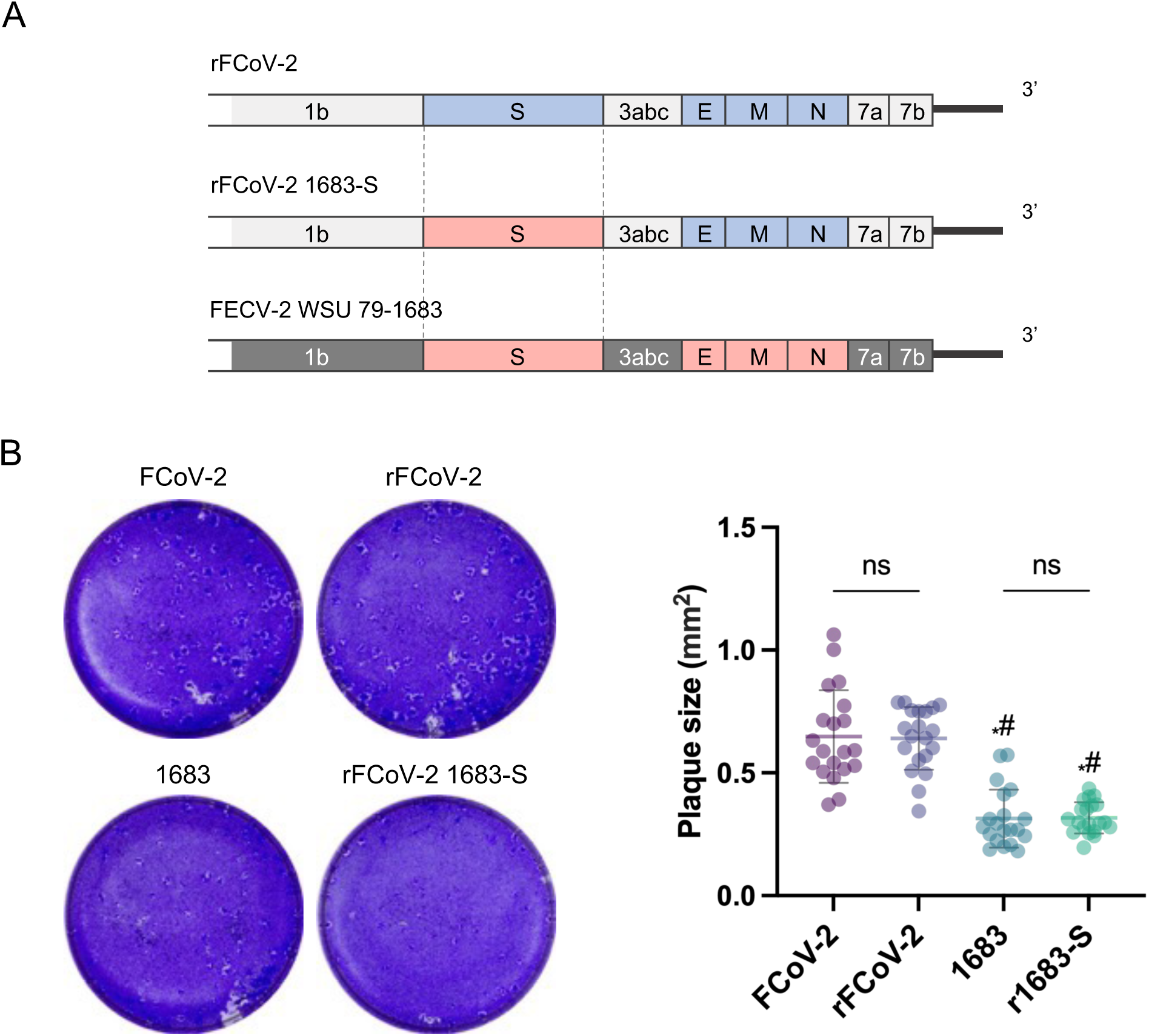
Construction of chimeric FCoV encoding spike gene from FECV-2. (A) Gene structures of parental viruses and chimeric FCoV encoding S gene from FECV-2. rFCoV-2 was used as a backbone for generating a chimeric FCoV expressing FECV S protein (1683-S) derived from FCoV-2 WSU 79-1683 strain, which is a serotype-2 FECV. (B) Plaque formation of parental FCoVs and rFCoVs in Fcwf-4 cells. Plaque assay was performed using Fcwf-4 cells inoculated with the parental FCoV-2, rFCoV-2, parental FECV (1683), and rFCoV-2 carrying 1683-S (r1683-S). Representative figures (left) and the sizes of plaques (n = 20 for each virus, right) are shown. Each dot in the graph indicates the diameter of a plaque. The mean ± SD was shown for each virus. Statistically significant differences versus parental FCoV-2 (*: *p* < 0.05) and recombinant FCoV-2 (#: *p* < 0.05) were determined by one-way ANOVA with Tukey’s test; ns, not significant.

### Construction of rFCoVs carrying a HiBiT gene

To prove the concept of generating recombinant reporter viruses using CPER, we constructed rFCoVs harboring NanoLuc-derived HiBiT tag in the N terminus of the ORF7b gene (Fig. 3A) by site-directed overlap extension PCR using the F10 plasmid. A fragment F10 possessing the HiBiT gene was amplified by PCR and subjected to CPER with the other fragments (F1-9 and the UTR linker). CPE was observed in CPER product-transfected cells, and the culture supernatants were collected at 4 dpt for rFCoV-1 HiBiT and 3 dpt for rFCoV-2 HiBiT. Then, each supernatant was subjected to passage in naïve Fcwf-4 cells followed by harvesting the P1 rFCoV-1 HiBiT and rFCoV-2 HiBiT at 3 dpi and 1 dpi, respectively. Growth kinetics of rFCoVs HiBiT were comparable to those of the parental viruses (Fig. 3B). The luciferase activity in cells infected with the P1 viruses increased from 12 to 48 or 72 hours (Fig. 3C). Subsequently, recombinant viruses were passaged multiple times to examine the stability of HiBiT gene in the viruses. The HiBiT tag was stably expressed after at least 5 passages (Fig. 3D).

**Fig 3.**
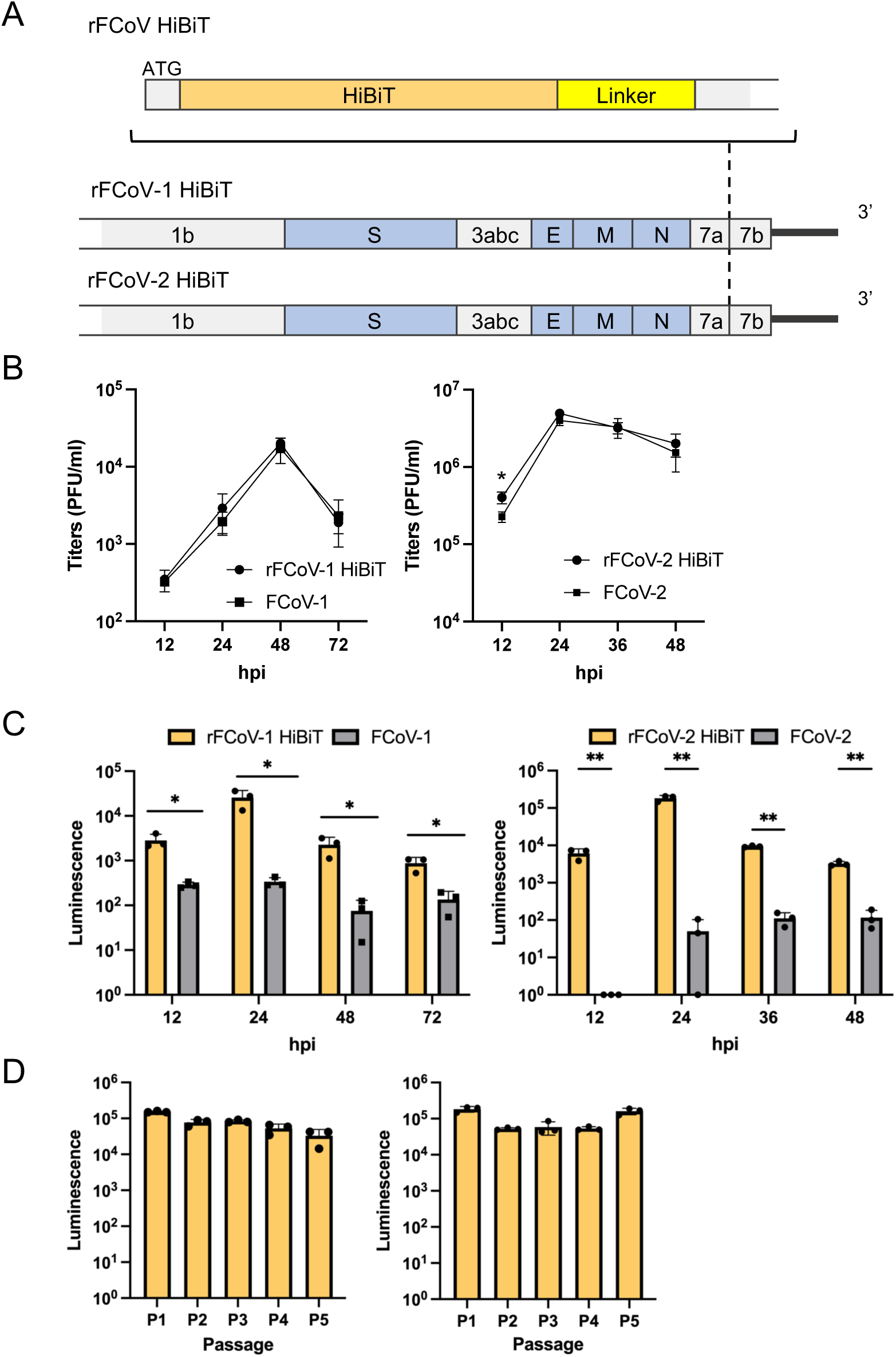
Construction of FCoV recombinants carrying a HiBiT gene. (A) Gene structure of FCoV recombinants carrying the HiBiT gene (rFCoV HiBiT). HiBiT gene was inserted immediately after the start codon (ATG) of the ORF7b followed by a linker. (B) Growth kinetics of rFCoVs HiBiT and parental FCoVs. Fcwf-4 cells were infected with the viruses at an MOI of 0.01, and the virus titers in the supernatants were measured from 12 to 48 or 72 hours post-infection (hpi). (C) Luciferase activities in Fcwf-4 cells infected with rFCoVs HiBiT and parental FCoVs. Cells were lysed followed by the addition of LgBiT, and the luciferase activities were determined from 12 to 48 or 72 hpi. (D) Stable expression of HiBiT tag during passage recombinant viruses. rFCoVs HiBiT were passaged five times (P1-P5) in Fcwf-4 cells. Luciferase activities in cells during each passage were determined at 24 hpi. In (B)-(D), the presented data were expressed as mean ± SD of triplicate samples. In (B) and (C), \**p* < 0.05, \*\**p* < 0.01 by a two-tailed Student’s *t* test without adjustment for multiple comparisons.

### Application of rFCoVs expressing HiBiT tag for assessment of antiviral inhibitors

Antiviral drugs, such as GS-441524, a nucleoside analog, and EIDD-1931, a viral RNA-dependent RNA polymerase (RdRP) inhibitor, have been applied in clinical settings for FIP (40, 41). To prove the reporter virus is appliable for examining antiviral inhibitors, Fcwf-4 cells infected with rFCoVs at an MOI of 0.01 for rFCoV-1 HiBiT and an MOI of 0.001 for rFCoV-2 HiBiT were respectively treated with GS-441524 (Fig. 4A) and EIDD-1931 (Fig. 4B) at the time of infection. The relative luciferase activity was determined at 24 hours for rFCoV-1 HiBiT and 18 hours for rFCoV-2 HiBiT. The luciferase activity was reduced in a dose-dependent manner. Half-maximal inhibitory concentrations (IC_50_) were 0.69 µM for GS-441524, 0.15 µM for EIDD-1931 against rFCoV-1-HiBiT, 0.69 µM for GS-441524, 0.066 µM for EIDD-1931 against rFCoV-2 HiBiT (Fig. 4A&B; Table 1) and no cytotoxicity was observed in the cells at the examined concentrations of all inhibitors (Fig. 4C). These data indicate that rFCoVs expressing HiBiT tag generated by CPER are useful for the assessment of antiviral inhibitors.

**Fig 4.**
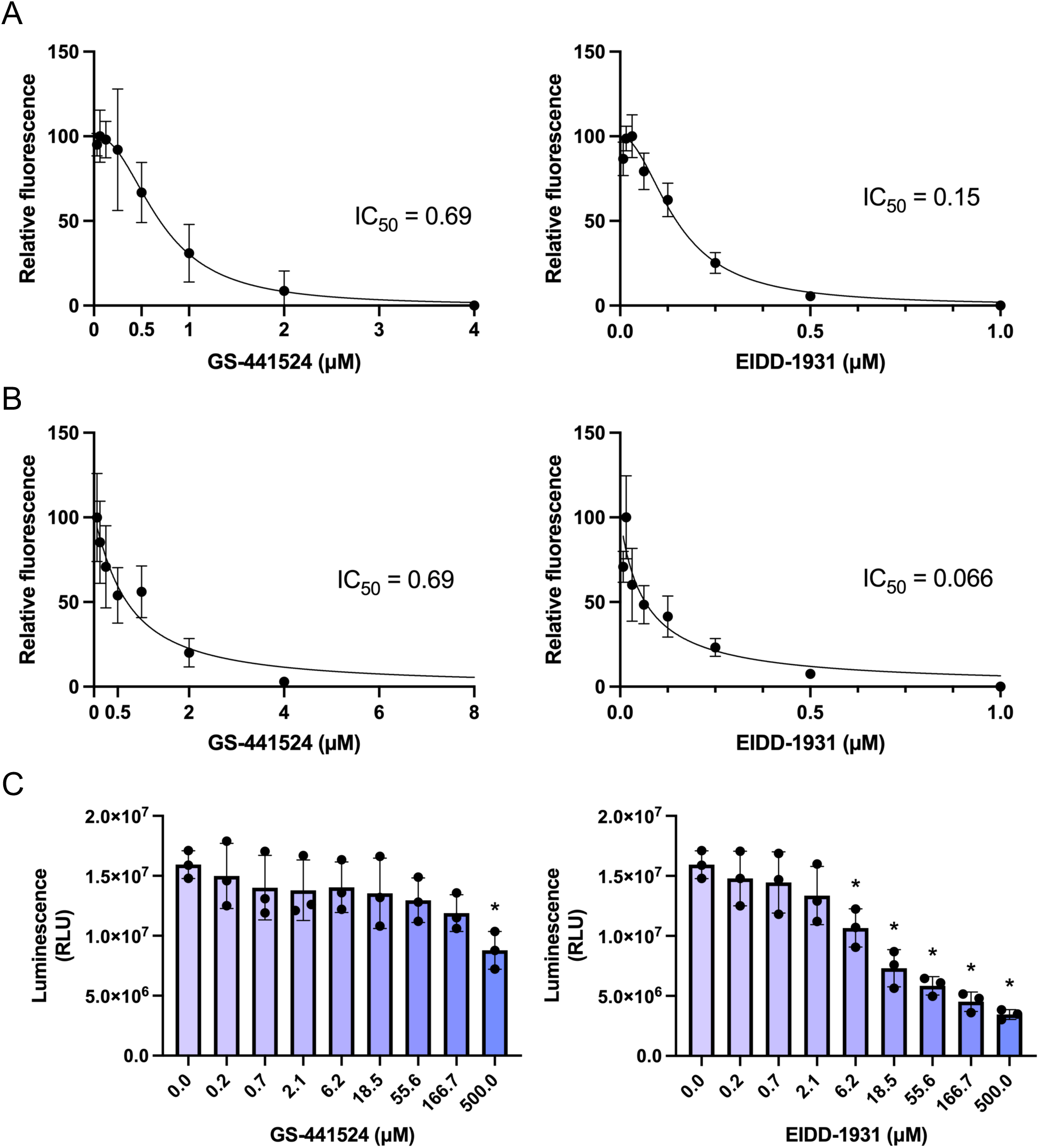
Application of reporter FCoVs for assessment of antiviral inhibitors. (A) IC_50_ of antiviral inhibitors against recombinant FCoV-1 carrying the HiBiT gene (rFCoV-1 HiBiT). Fcwf-4 cells infected with rFCoV-1 HiBiT at an MOI of 0.01 were treated with GS-441524 and EIDD-1931 at the time of infection. The luciferase activity levels were determined by a luciferase assay at 24 hours. Relative fluorescence was calculated by intensity of luciferase activity in the cells treated with antiviral inhibitors in comparison to mock control. (B) IC_50_ of antiviral inhibitors against recombinant FCoV-2 carrying the HiBiT gene (rFCoV-2 HiBiT). Fcwf-4 cells infected with rFCoV-2 HiBiT at an MOI of 0.001 were treated with GS-441524 and EIDD-1931 at the time of infection. The luciferase activity levels were determined by a luciferase assay at 18 hours. Relative fluorescence was calculated by intensity of luciferase activity in the cells treated with antiviral inhibitors in comparison to mock control. (C) Cytotoxicity of antiviral inhibitors. Fcwf-4 cells were treated with GS-441524 or EIDD-1931 for 24 hours. Cell toxicity was evaluated by measurement of ATP in live cells. The presented data were expressed as mean ± SD of triplicate samples. Statistically significant differences versus cell control (\**p*< 0.05) were determined by one-way ANOVA with Dunnett’s test.

**Table 1.** IC_50_ of antiviral inhibitors determined by luciferase assay.

| Inhibitor | IC<sub>50</sub> (μM) |  |
| --- | --- | --- |
|  | rFCoV-1 HiBiT | rFCoV-2 HiBiT |
| GS-441524 | 0.69 | 0.69 |
| EIDD-1931 | 0.15 | 0.066 |

## Discussion

Herein, we successfully established the rapid and versatile reverse system for FCoVs. Using the CPER method, we demonstrated the capability of the recombinant viruses to be introduced desired modifications or exogenous short reporter genes without any replication deficiency and genetic instability. CPER allows us to generate a full-length cDNA of FCoV with a simple and rapid PCR-based method from plasmids carrying viral fragments and to easily introduce modifications within a couple of days. To construct a full-length clone with desired mutations usually requires several days with the previously reported methods, and therefore, our system will accelerate reverse genetics studies of FCoVs.

In this study, we applied the CPER method to generate chimeric FIPV-2 encoding FECV-2 S gene. We successfully rescued the chimeric virus expressing S protein derived from the same serotype virus and showed switching of the phenotypes depending on S protein. While further studies are needed for chimeric viruses based on rFCoV-1 and also various combinations, such as chimeras of ORF3c and/or ORF7b, we have proved that our system allows to modify FCoV genes easily. The development of recombinant viruses for both serotypes and their chimeras will contribute to further analysis and comparison of the characteristics between serotypes.

The CPER-based method has been used for studying a variety of mutations of SARS-CoV-2, especially on S gene, during the COVID-19 pandemic to understand the molecular mechanisms of pathogenesis (31, 39, 42–44). Due to the difficulty in isolating FCoV, it can be challenging to immediately examine the causative virus, particularly its S protein, a major determinant of virus characteristics including pathogenicity, during FIP outbreaks. The recent outbreak of FIP in Cyprus was caused by FCoV-2 (referred to as FCoV-23), which exhibits high pathogenicity and transmissibility from cats to cats (9). The risk of spreading the outbreak is a concern, as evidenced by the imported case in the UK (45). Despite the significant impact on the cat population, there have been limited reports to examine the virological features of this highly pathogenic FCoV (46). For the quick development of countermeasures against emerging FCoVs such as FCoV-23, CPER method will be useful in the early stage of an outbreak without access to virus samples.

The antivirals for FIP treatment have been becoming available in some parts of the world, but concerns remain regarding the potential emergence of drug resistant FCoVs following treatment. Therefore, the development of effective therapeutics with different mechanisms for FIP is still needed. In the present study, we demonstrated recombinant FCoVs carrying HiBiT gene, showing IC_50_ values consistent with previous studies (47, 48). Since the luciferase assay examining the HiBiT tag expression in cells could quantify viral load within 30 min with enough sensitivity without virus titrations and RNA extractions, our system is useful for assessing antiviral inhibitors. While there can be a limitation to the insertion of a longer reporter gene, which reduced viral replication (data not shown), our HiBiT reporter viruses may be applicable for a wide variety of experiments that require the sensitive quantification of virus replication. The identification of a suitable gene locus for the insertion of the longer gene requires further investigation.

In summary, we established an efficient reverse genetics system for both FCoV-1 and -2 that can be beneficial for several molecular virological applications, because the system allows us to rapidly introduce desired modifications and reporter genes into FCoV genome. Thus, our system will facilitate investigations into the molecular mechanisms of pathotype switch of FCoV and also for understanding viral life cycles and screening of antiviral drugs.

## Materials and Methods

### Cells

*Felis catus* whole fetus 4 (Fcwf-4) cells and human embryonic kidney 293T (HEK293T) cells were maintained with 5% CO_2_ at 37 °C in high-glucose Dulbecco’s modified Eagle’s medium (DMEM; Nacalai Tesque) supplemented with 10% fetal bovine serum (FBS; Nichirei Bioscience), 100 U/ml penicillin, and 100 μg/ml streptomycin (Gibco).

### Viruses

Serotype-1 FCoV strain C3663 was initially isolated from a cat with FIP in 1994 (33) and propagated in Fcwf-4 cells as described previously (7). Serotype-2 FCoV strain WSU 79-1146 (34, 35) and WSU 79-1683 (37) obtained from ATCC were used. To prepare the viral stock, the seed virus was inoculated into monolayered Fcwf-4 cells and cultured until CPE was confirmed. The cultured medium was collected and centrifuged, and then the supernatants were stored as a viral stock at −80°C until use.

### Viral genome sequencing

The virus sequences were verified by viral RNA-sequencing analysis. Viral RNA was extracted from the supernatant of infected cells using NucleoSpin RNA Virus (TaKaRa Bio). The sequencing library for total RNA sequencing was prepared using the KAPA RNA HyperPrep Kit for Illumina (Roche) or Collibri Stranded RNA Library Prep Kit for Illumina systems (Thermo Fisher Scientific). Paired-end, 300 bp sequencing was performed using MiSeq (Illumina) with MiSeq reagent kit v3 (Illumina) or NextSeq 2000 (Illumina) with NextSeq 1000/2000 P1 XLEAP-SBS Reagent Kit (Illumina). Sequencing reads were trimmed and mapped to reference sequences using CLC Genomics Workbench v22.0.2 (Qiagen).

### cDNA synthesis

First strand cDNA was synthesized by using PrimeScript Ⅱ 1st stranded cDNA Synthesis Kit (TaKaRa Bio) with random hexamer primers according to the manufacturer’s protocol.

### Plasmid construction

FCoV genome was divided into 10 fragments (F1-F10) which are up to 4,672 base pairs in length and cover the entire FCoV sequence, were amplified from the cDNA using specific primer sets for each serotype of FCoV and PrimeSTAR GXL DNA polymerase (TaKaRa Bio). Each fragment was cloned into the pGEM-T Easy vector. A UTR linker for FCoV were generated using the pUCFa vector which encodes sequences of 18 nt of the 3’UTR of FCoV, HDVr, BGH polyA signal, CMV promotor and 70 nt of the 5’UTR of FCoV. Two silent mutations (i.e. strain C3663; C6444A and A6447C and strain WSU 79-1146; A6856G and C6859T) were introduced into the F3 plasmid respectively as a genetic marker by site-directed overlap extension PCR. The construct of HiBiT-tagged FCoV was prepared by introducing HiBiT tag (VSGWRLFKKIS) and a linker (GSSG) sequence into the 5’ terminus of the ORF7b gene. To construct chimeric FCoV-2 encoding S gene from strain WSU 79-1683, the cDNA of the S was replaced into the F8 plasmid of FCoV-2. Nucleotide sequences of all constructs were confirmed by sanger sequencing or whole plasmid sequencing service (Azenta). Then, DNA fragments were amplified with specific primers for subsequent CPER assembly.

### CPER assembly

Recombinant FCoVs were generated by CPER as described previously (Torii et al., 2021) with some modifications. The plasmids encoding FCoV gene fragments (F1-F10) and the UTR linker were used to generate viral cDNA fragments. PCR fragments, having complementary ends with at least 13 nt overlap for CPER, were amplified with specific primers. The purified 11 fragments (F1 to F10 and the UTR linker) were mixed in equimolar amounts (0.1 pmol each) in a 50µl reaction volume containing 2 µl of PrimeSTAR GXL DNA polymerase and subjected to CPER. The cycling condition of CPER was as follows: an initial 2 minutes of denaturation at 98°C, 35 cycles of 10 s at 98°C and 15 minutes at 68°C, and a final extension for 15 minutes at 68°C.

### Recovery of rFCoV

The CPER products without purification were transfected into Fcwf-4 cells co-cultured with HEK293T cells with Trans IT LT-1 (Mirus) in accordance with the manufacturer’s instructions. Viral recovery was confirmed by CPE and the supernatant was passaged in Fcwf-4 cells to produce P1 virus for conducting experiments.

### Indirect immunofluorescence assay

Fcwf-4 cells inoculated CPER generated virus were fixed with 4% paraformaldehyde when CPE was observed, and permeabilized for 10 min at room temperature with PBS containing 0.1% Triton X-100. The cells were stained with Coronavirus pan Monoclonal Antibody FIPV3-70 (Thermo Scientific) and Goat anti-Mouse IgG (H+L) Alexa Fluor 488 (Thermo Scientific) and then mounted with 4’,6-diamidino-2-phenylindole (DAPI; Dojindo). Immunofluorescence microscopy was performed with a Zeiss Axio Vert.A1 Inverted Microscope for Phase Contrast.

### Titration

The infectious titer in the culture supernatants were determined by the plaque assay. The culture supernatants were inoculated onto Fcwf-4 cells in 12-well plates after ten-fold serial dilution for 1h at 37°C.After washing with DMEM, a mounting solution containing 0.8% agar (FUJIFILM WAKO) or 1% methylcellulose (Sigma-Aldrich) in Eagle’s MEM (EMEM; Nissui Pharmaceutical Co) supplemented with 2% FBS was overlaid onto the cells. The cells were incubated at 37°C and fixed at 1 or 2 dpi with 10% Formalin Neutral Buffer Solution, Deodorized (FUJIFILM WAKO) and stained with crystal violet (Sigma-Aldrich).

### Growth kinetics

The culture supernatants were inoculated onto Fcwf-4 cells in 6-well plates at an MOI of 0.01. After 1 h of incubation, cells were washed once and cultured in DMEM with 2% FBS. The culture supernatants were harvested at 12, 24, 48, 72 h for FCoV-1 and at 12, 24, 36, 48 h for FCoV-2, after inoculation. Virus titers were evaluated by plaque assay.

### HiBiT luciferase assay

Luciferase activity was measured with GloMax Discover Microplate Reader (Promega) using a Nano-Glo HiBiT Lytic assay system (Promega) according to the manufacturer’s protocol. In brief, Fcwf-4 cells seeded on 96-well plate were infected with rFCoVs HiBiT at an MOI of 0.01. The Nano-Glo HiBiT Lytic reagent was prepared by diluting the LgBiT protein 1:100 and the substrate solution 1:50 into an appropriate volume of lytic buffer. The reagent was added to the cells infected with viruses and mixed well, followed by 10 minutes of incubation. The luciferase activity was measured following equilibration of the reactants. The measured luminescence intensity values were designated as relative light units (RLU).

### Inhibitors

GS-441524 and EIDD-1931 were purchased from Selleck Chemicals, dissolved in dimethyl sulfoxide (DMSO) and stored at 100 mM at −30°C until use.

### Inhibitor tests with rFCoVs expressing HiBiT tag

Antiviral inhibitors were serially diluted in 2-fold increments by DMEM containing 2% FBS and plated on 96-well microplates. The diluted inhibitors in the plates were mixed with Fcwf-4 cell suspensions and rFCoV-1 HiBiT or rFCoV-2 HiBiT at an MOI of 0.01 and 0.001, respectively. After 24 hours culture for FCoV-1 and 18 hours culture for FCoV-2, the luciferase activity was measured. IC_50_ values were defined in GraphPad Prism version 10.2.3 (GraphPad Software) with a variable slope. Nontreated cells were used as a control for 100%.

### Compound cytotoxicity assay

Cells were treated with GS-441524 or EIDD-1931 diluted in 3-fold serial increments by DMEM containing 2% FBS. After 24 hours culture, cell toxicity was evaluated by measurement of ATP in live cells with CellTiter-Glo 2.0 reagent (Promega). Luminescence was detected with GloMax Discover Microplate Reader (Promega).

### Statistical analysis

The data were expressed as mean ± SD. Statistical significance was tested using two-tailed Student’s *t* test (Fig. 1D, 3B and 3C), one-way ANOVA with Tukey’s test (Fig. 2B), one-way ANOVA with Dunnett’s test (Fig. 4C). All statistical tests were performed using GraphPad Prism version 10.2.3 (GraphPad Software).

## Data Availability

This study did not generate any unique datasets.

## Acknowledgement

We thank Dr. Michihito Sasaki (International Institute for Zoonosis Control, Hokkaido University, Japan), Dr. Koshiro Tabata (Institute for Vaccine Research and Development, Hokkaido University, Japan) and Asako Shigeno (International Institute for Zoonosis Control, Hokkaido University, Japan) for their assistance. This work is supported by JSPS KAKENHI Fund for the Promotion of Joint International Research (International Leading Research) (JP23K20041 to Izumi Kida and Keita Matsuno), KAKENHI Grant-in-Aid for Scientific Research B (21H02736 to Takasuke Fukuhara), AMED ASPIRE (JP23jf0126002 to Izumi Kida and Keita Matsuno), AMED CREST (JP22gm1610008 to Takasuke Fukuhara), JST SPRING (JPMJSP2119 to Izumi Kida), Hokuto Foundation for Bioscience (to Tomokazu Tamura) and Grants-in-Aid for R&D of Young Scientist from Northern Advancement Center for Science &Technology (NOASTEC) of Hokkaido (to Tomokazu Tamura).

